# Synthesis of the NarP response regulator of nitrate respiration in *Escherichia coli* is regulated at multiple levels by Hfq and small RNAs

**DOI:** 10.1101/2021.03.17.435884

**Authors:** Anaïs Brosse, Anne Walburger, Axel Magalon, Maude Guillier

**Author notes:** Present address : Université Paris-Saclay, INRAE, AgroParisTech, Micalis Institute, 78350 Jouy-en-Josas, France (Anaïs Brosse).

## Abstract

Two-component systems (TCS) and small RNAs (sRNA) are widespread regulators that participate in the response and the adaptation of bacteria to their environments. They mostly act at the transcriptional and post-transcriptional levels, respectively, and can be found integrated in regulatory circuits, where TCSs control sRNAs transcription and/or sRNAs post-transcriptionally regulate TCSs synthesis.

In response to nitrate and nitrite, the paralogous NarQ-NarP and NarX-NarL TCSs regulate the expression of genes involved in anaerobic respiration of these alternative electron acceptors. In addition to the previously reported repression of NarP synthesis by the SdsN_137_ sRNA, we show here that RprA, another Hfq-dependent sRNA, also negatively controls *narP*. Interestingly, the repression of *narP* by RprA actually relies on two independent controls. The first is *via* the direct pairing of the central region of RprA to the *narP* translation initiation region and presumably occurs at the translation initiation level. In contrast, the second control requires only the very 5’ end of the *narP* mRNA, which is targeted, most likely indirectly, by the full-length or the shorter, processed, form of RprA. In addition, our results raise the possibility of a direct role of Hfq in *narP* control, further illustrating the diversity of post-transcriptional regulation mechanisms in the synthesis of TCSs.

## INTRODUCTION

Bacteria have the ability to sense, respond and adapt to a wide diversity of environments and their capacity to regulate gene expression plays a key role in this process. Examples of control have been reported at multiple steps of gene expression. Transcription initiation is, for instance, commonly regulated by proteins that bind to the promoter regions of genes and can activate or repress their transcription [1]. Regulators falling in this category are involved in the response to diverse input signals, often via a change in their activity in response to a cognate signal, by phosphorylation in the case of two-component systems (TCS). TCSs are widely used in bacteria and consist of a sensor kinase that can auto-phosphorylate in response to specific stimuli and transfer the phosphate group to a cognate response regulator (RR). In most cases, the phosphorylated form of the RR is the biologically active form and regulates transcription by binding to DNA.

While undoubtedly beneficial for bacterial adaptation, transcriptional regulation is not the only form of control. Indeed, many bacterial genes can also be regulated at the post-transcriptional level and, in these cases, translation is often the regulated step. Although translational control by proteins was described many decades ago, the observation that a myriad of small RNAs, most of which acting as post-transcriptional regulators, exist in virtually all bacteria has confirmed the importance of post-transcriptional control in bacterial adaptation. In the vast majority of cases, sRNAs act by pairing to target-mRNAs via imperfect base-pairing interactions and repress, or more rarely increase, their translation and/or stability [2]. Based on the examples studied so far, the most common scenario is that sRNAs of this category pair at or in the vicinity of the ribosome binding-site of their target and repress translation initiation by directly competing with binding of the 30S ribosomal subunit. Several other mechanisms have been described, however, including translation activation or repression by sRNAs pairing outside of the translation initiation region (TIR), or the stabilization or destabilization of target-mRNAs as a direct consequence of sRNAs binding [3,4]. In enteric bacteria such as *E. coli* and *Salmonella*, for which many of the details of sRNAs action have been elucidated so far, these imperfectly pairing sRNAs require an RNA chaperone, Hfq or the more recently identified ProQ, for stability and duplex formation with their targets [5,6]. Consistent with this, Hfq is involved in the regulation of a multitude of genes whose expression is under the control of sRNAs. Furthermore, Hfq has also been shown to be involved in the direct control of gene expression, independently of sRNAs [7,8].

Interestingly, transcriptional and post-transcriptional controls do not form completely independent regulatory networks in bacterial cells but rather result in mixed regulatory circuits relying on both proteins and sRNAs acting mostly at the transcriptional and post-transcriptional level, respectively [9,10]. Transcription of sRNAs is most often controlled by transcriptional regulators, while sRNAs in turn post-transcriptionally regulate the synthesis of transcriptional regulators. One example is the stress response alternative sigma factor RpoS, which directs transcription of SdsR and SdsN sRNAs in enterobacteria [11,12], while RpoS synthesis is directly up-regulated by at least three sRNAs, namely DsrA, RprA and ArcZ [13–16]. This control of, and by, sRNAs is also true for regulators of TCSs, including OmpR and PhoP, two of the most studied response regulators in enterobacteria, as well as the LuxO RR involved in quorum-sensing in Vibrio species [17–23].

NarP is another example of a RR whose synthesis is under sRNA control. Together with its paralog NarX-NarL, the NarQ-NarP TCS regulates the expression of genes involved in the anaerobic respiration on nitrate, the energetically most favourable electron acceptor in the absence of oxygen, and on nitrite, the reduction product of nitrate [24]. The translation of *narP* mRNA is repressed by SdsN_137_, one isoform of a set of RpoS-dependent sRNAs [12]. In addition to this control, we report here that expression of *narP* is also regulated by Hfq and the RprA sRNA, whose synthesis is primarily controlled by the Rcs phosphorelay, but also responds to the CpxAR TCS and the LrhA regulator of flagellar synthesis [15,25,26]. RprA regulates expression of multiple genes involved in diverse stress responses, biofilm formation, formate metabolism, conjugation as well as the gene encoding LrhA, one of its transcriptional regulator [14,27–29]. By adding *narP* to the list of RprA and Hfq targets, our results expand the connections between sRNAs and TCSs, two classes of widespread bacterial regulators, and highlights the high level of integration of the diverse pathways that regulate gene expression. In addition, they demonstrate a dual level of control by a single sRNA, acting both canonically at the translation initiation region (TIR) of *narP* and, via a distinct sRNA site, and most likely indirectly, at the distant 5’end of the same mRNA.

## MATERIALS AND METHODS

### Strains, plasmids and general microbiology techniques

All strains and plasmids used in this study are listed in Supplementary Table S1. Cells were grown at 37°C, either in LB medium or in a defined medium (MMGly) composed of potassium phosphate buffer (100 mM) adjusted to pH 7.4, ammonium sulfate (15 mM), sodium chloride (9 mM), magnesium sulfate (2 mM), sodium molybdate (5 µM), Mohr’s salt (10 µM), calcium chloride (100 µM), casaminoacids (0.5 %) and thiamine (0.01 %), supplemented with 140 mM of glycerol as sole carbon source. When needed, nitrate was added at a final concentration of 5 mM. Anaerobic growth was performed in gas tight Hungate tubes under Argon atmosphere. When necessary, antibiotics were used at the following concentrations: 100 or 150µg/ml ampicillin (Amp_100_ or Amp_150_ in figure legends), 10µg/ml tetracycline, 25μg/ml or 50µg/ml kanamycin and 10µg/ml chloramphenicol. 100 µM of IPTG was also added when required to induce expression of sRNAs from pBRplac derivatives. Amplification of DNA fragments was performed with either Phusion DNA polymerase or LongAmp DNA polymerase (NEB).

Except for cloning, strains are all derivatives of *E. coli* MG1655, modified by recombineering or P1 transduction when needed. Mutant *rprA::tet* (from strain NM667, from N. Majdalani, unpublished) and *sdsN::kan* (strain GS0762 [12]) were obtained from S. Gottesman’s and G. Storz’s laboratories, respectively, while the *ΔnarP::kan* allele was taken from the Keio collection (strain JW2181, [81]). For the construction of strains carrying *hfq* point mutants, a *Δhfq::cat-sacB purA::kan* mutant (strain DJS2604, from D. Schu) was first transduced into the recipient strain, and the different *hfq* alleles (from strains DJS2927 (wt), DJS2609 (ΔHfq), KK2561 (R16A), KK2562 (Y25D) or KK2560 (Q8A), from D. Schu carrying alleles from [31,54]) were then moved into the resulting strain, allowing selection on glucose minimal medium. The various fusions with the *lacZ* reporter gene were made as follows: a PCR fragment encompassing the sequence to be placed upstream of *lacZ*, flanked by homology regions, was recombined into the strain MG1508 (or MG2114 for construction of the P_napF_-lacZ fusion), carrying the genes for recombineering on a mini-lambda and where a P_tet_-*cat-sacB* cassette has been introduced upstream of *lacZ*. Recombinant cells were selected on sucrose-containing medium, checked for chloramphenicol sensitivity and the fusion was sequenced. For fusions whose transcription originates from P_narP_ or P_napF_, an *rrnBt2* transcription terminator was introduced upstream, either on the PCR fragment (P_narP_), or by recombineering into MG2114 strain carrying an *rrnBt2*-*cat-sacB-lacZ* construct (for P_napF_ promoter fusion). Fusions P_narP-207+50_-*lacZ* and P_tet+1+50narP_-*lacZ* have a mutation in the second codon of *lacZ* (ACC is changed to AAC) which prevents the formation of an inhibitory structure for translation.

For the insertion of a 3x-Flag (sequence DYKDHDGDYKDHDIDYKDDDDK) just upstream of the *narP* stop codon, a PCR fragment carrying the 3x-Flag sequence preceded by a linker (protein sequence GAGAGAGA) and followed by a kanamycin resistance gene flanked by FRT sites was amplified from plasmid pSUB11 [82] with homology regions to the end of *narP*. This fragment was then recombined into strain MG1433 and checked by sequencing after selection and purification on LB-Kan. A control adding only the linker sequence was constructed as well. This *narP-*3xFlag -FRT-KanR-FRT allele was then transduced into recipient strains as needed.

The plasmids used to overexpress the different Hfq-dependent sRNAs are mostly from (Mandin & Gottesman, 2010), with the addition of pCsrB (from N. de Lay, unpublished), pMcaS, pMicL and pSdsN_137_ (from G. Storz, [12,38,39]) and pCpxQ and pDapZ (this study, based on [40,41]; see Fig. S1 for Northern-blot validation of their overexpression). Mutant plasmid pSdsN_137_-1 [12] was obtained from G. Storz and pRprAmut2 was constructed by amplification of the pRprA plasmid with mutagenic primers using the Pfu enzyme (Agilent), followed by DpnI digestion and transformation into the cloning strain NEB5-alpha F’I^q^. Table S2 summarizes the main oligonucleotides used in this study to construct strains or plasmids.

### Measure of β-galactosidase activity

Cells were diluted 250-fold into fresh medium from an overnight culture and grown to mid-exponential phase and β-galactosidase activity was measured following Miller’s protocol [83]. Cells were lysed either with toluene (for aerobic extracts) or SDS-chloroform (anaerobic extracts). Results presented here correspond to the average of at least two independent experiments and error bars correspond to the standard deviation.

### RNA extraction and northern-blot analysis

The RNAs were extracted with hot phenol as previously described [19] from the same cultures as those used for β-galactosidase assays. A constant amount of total RNA (between 3.5 and 14 μg) was loaded on 8% urea acrylamide gel in 1X TBE and transferred to a nitrocellulose membrane (GE Healthcare). For detection, we used specific 5 ’ end biotinylated probes (see Table S2 for sequence), hybridized to the membrane overnight in Ultrahyb buffer (Ambion). After two washes in SSC 2X-SDS 0.1% and two washes in SSC 0.1X–SDS 0.1% at 42°C, the membrane was incubated in nucleic acid detection blocking reagent (ThermoScientific), and then in the same solution in presence of a streptavidin-alkaline phosphatase conjugate (Life Technologies). Membrane was then washed three times in wash buffer (Na_2_HPO_4_ 29mM, NaH_2_PO_4_ 8.5mM, NaCl 34mM and SDS 0.05%), equilibrated in assay buffer (NaCl 0.1M, Tris 0.1M pH 9.5) and chemiluminescence was detected using the CDP-star substrate (Applied Biosystems).

### Western blot and Phos-tag electrophoresis

The whole procedure for protein extraction, gel electrophoresis and western-blot detection from Phos-tag containing gels was performed as described previously in [17], except that precast gels containing Phos-tag and Zn^2+^ (Wako) were used. For standard western blots, proteins were separated on precast TGX gels (Biorad). A monoclonal anti-Flag antibody conjugated to alkaline phosphatase (Sigma) was used for the detection of the flagged proteins, with CDP-star reagent to detect chemiluminescence.

## RESULTS

### Hfq controls the expression of several *Escherichia coli* response regulators

As mentioned above, several examples of genes encoding regulators of TCSs whose expression is post-transcriptionally regulated by Hfq-dependent sRNAs have been reported. TCSs are widespread in bacteria and more than 30 RR genes exist in the model bacterium *E. coli* for instance. This raises the possibility that other RRs could be subject to sRNA control. We have begun to address this question by using *lacZ* reporter fusions to follow the production of 11 *E. coli* regulators of TCSs (listed in Table 1). RRs such as OmpR, PhoP or NarP, whose control by sRNAs has been previously described, have been included in this set. It is worth noting that, in agreement with their control by sRNAs, their mRNAs were found deregulated in an *hfq* null strain and/or enriched following Hfq co-immunoprecipitation [30– 32]. The other RRs included in this study set were BaeR, EvgA, HprR, KdpE, RstA and UvrY, for which various indications of sRNA control have been reported, as well as BasR and PhoB, for which no indication for sRNA control is available to our knowledge (Table 1).

**Table 1:** TCS investigated in this study.

| Two-component system<br>(Kinase-Regulator)<sup>1</sup> | Number of genes<br>whose expression is<br>up/down in the TCS<br>mutant<br>(based on [33]) | Indication for sRNA regulation<sup>4</sup> |  | Portion of the RR gene<br>present on the fusion<br>(relative to start codon)<sup>5</sup> |
| --- | --- | --- | --- | --- |
| | | Deregulation in $\Delta hfq$ | Interaction with Hfq | |
| BaeS-BaeR<sup>2</sup><br>(response to membrane stress) | 14/8 |  | ++ [30]<br>+ for 3' UTR [32] | -30 +720 |
| BasS-BasR<br>(response to iron) | 20/12 |  |  | -18 +666 |
| EvgS-EvgA<br>(antibiotic resistance and acid stress) | 2/3 |  | Slight enrichment upon Hfq IP, and presence in chimeric fragments with sRNAs [34,35] | -125 +612 |
| KdpD-KdpE<sup>2</sup><br>(potassium transport) | 10/10 | | ++ [30]<br>++ in $\Delta proQ$ [35] | -39 + 675 |
| NarQ-NarP<sup>3</sup><br>(nitrate metabolism) | NarQ 20/25<br>NarP 3/9 | +++ [30] | +++ [30,31]<br>+ [32] | -150 +645 |
| PhoR-PhoB<br>(phosphate metabolism) | 1/8 |  |  | -41 +687 |
| RstB-RstA<br>(acid stress response) | 12/22 |  | Presence in chimeric fragments with sRNAs [35] | -13 +726 |
| BarA-UvrY<br>(global metabolism) | BarA 2/24<br>UvrY 86/37 |  | +++ [30]<br>Slight enrichment upon Hfq IP, and presence in chimeric fragments with sRNAs [35] | -44 +654 |
| HprS-HprR<sup>3</sup><br>(response to oxidative stress) | 6/7 |  | ++ in only one out of two experiments [31] | +1 +669 |
| EnvZ-OmpR<br>(response to osmotic stress and acid stress) | 71/54 |  | + [31] | -35+30 |
| PhoQ-PhoP<br>(response to magnesium) | 4/27 | ++ [30,36] | +++ [30,31]<br>+ [32]<br>Slight enrichment upon Hfq IP, and presence in chimeric fragments with sRNAs [34,35] | -36+30 |

<sup>1</sup>unless otherwise indicated, TCSs are expressed from a bicistronic operon with the gene for the response regulator being the first cistron.
<sup>2</sup>In the BaeS-BaeR and KdpD-KdpE systems the kinase is the first gene in the operon
<sup>3</sup>The two genes of the NarQ-NarP TCS are encoded by different loci of the genome.
<sup>4</sup>Weak, moderate or strong indications for sRNA regulation in different studies are indicated by +, ++ or +++, respectively.
<sup>5</sup>Positions of the transcription start sites (TSS) were chosen based on [37]; even though *baeR* and *kdpE* are not the first gene of their operon, they were introduced downstream of the P<sub>tet</sub> promoter in the absence of the *baeS* or *kdpD* gene, respectively. Note that the start codon is GUG for *kdpE* and *rstA*, and UUG for *uvrY*.

Translational fusions of *lacZ* to these 11 regulators were constructed at the *lacZ* locus by recombineering (Fig. 1A). All fusions followed the same general organization. First, their transcription was systematically driven by a constitutively expressed P_tet_ promoter to exclude regulation at the promoter level. Second, the sequence of each RR gene covering the 5’ UTR and all of the coding region with the exception of the stop codon was placed in frame upstream of *lacZ* starting at the 10^th^ aminoacid of β-galactosidase; for *ompR* and *phoP* however, only the 10 first aa of the coding sequence were included in the fusions as this was previously shown to be sufficient to allow regulation by sRNAs (13, 14). Expression of these different fusions was then measured in an *hfq*^*+*^ or *hfq*^*-*^ background, with the idea that a Hfq effect could indicate potential regulation by sRNAs. Note however that genes whose expression is not affected by Hfq in this experiment could nonetheless be regulated by sRNAs, for instance because these sRNAs are expressed in different experimental conditions than those used here or are Hfq-independent, or because the fusion is not a good reporter in these cases.

**Figure 1:**
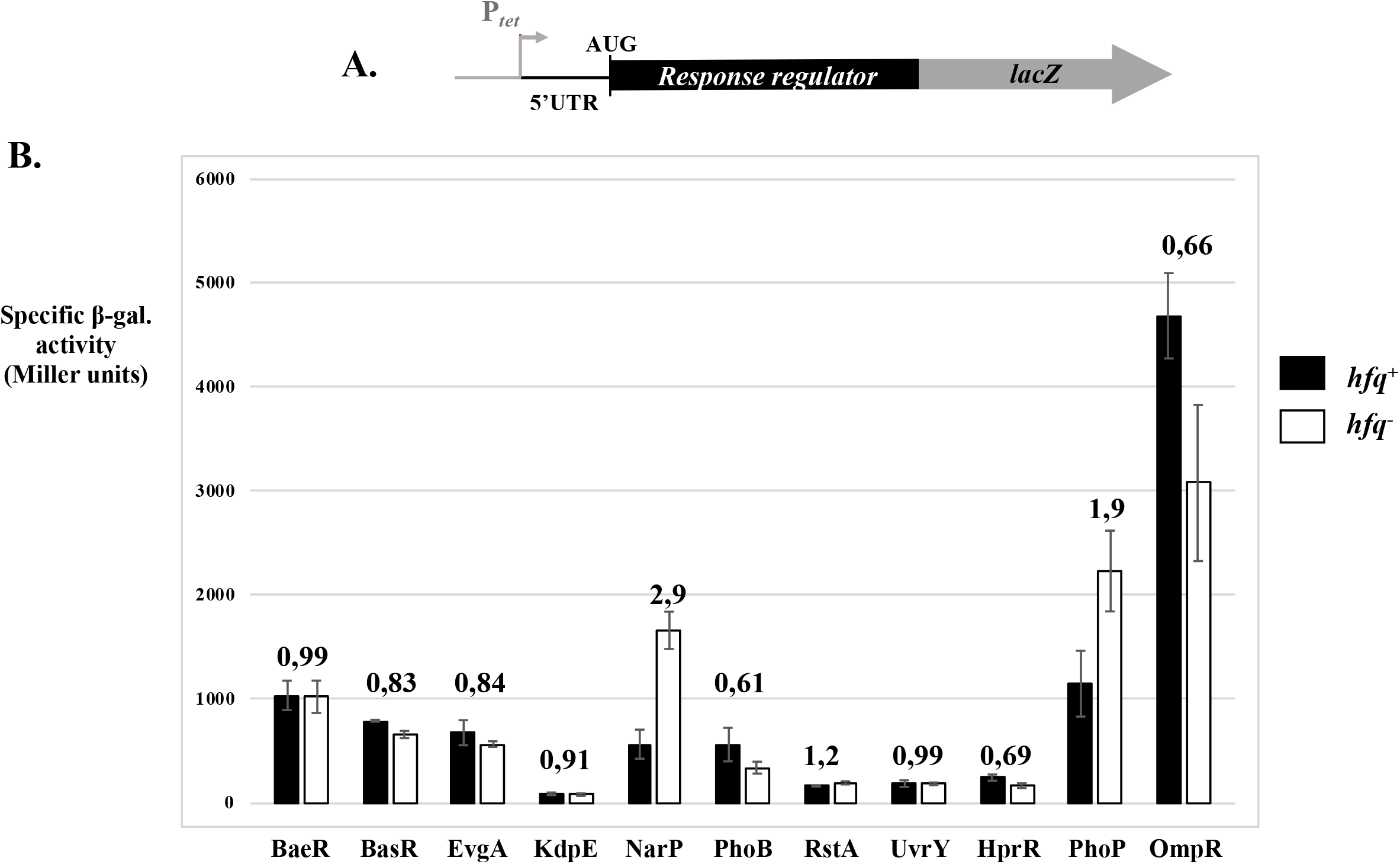
Hfq modulates the expression of several RR genes. (A) Scheme of the different translational fusions used to follow RR expression. All fusions are expressed from a P_tet_ promoter; *phoP* and *ompR* coding regions present on the fusions are limited to the 10 first aminoacids, while the entire coding regions of the RR genes, except the stop codon, are present in all other fusions. (B) The β-galactosidase activities of each fusion were measured in an *hfq*^+^ of *hfq*^-^ background in exponential phase in LB medium. wt and *hfq* null strains used here were, respectively, AB1000 and AB1009 (*baeR*); AB1001 and AB1010 (*basR);* AB1002 and AB1011 (*evgA*); AB1003 and AB1012 (*kdpE*); AB1004 and AB1013 (*narP*); AB1005 and AB1014 (*phoB*); AB1006 and AB1015 (*rstA*); AB1007 and AB1016 (*uvrY*); AB1008 and AB1017 (*hprR*); MG1511 and MG1515 (*phoP*) and AB1148 and AB1149 (*ompR*).

Consistent with previous data [36], expression of the *phoP* fusion was up-regulated almost 2-fold in the *hfq* mutant, even if it is unclear at this stage whether this is due to sRNA control or not. In contrast, expression of *ompR*, also subject to negative regulation by sRNAs, namely OmrA and OmrB (14), was down-regulated 1.5-fold in the absence of Hfq, possibly indicating the existence of sRNAs that can activate *ompR* expression. Expression of *baeR, basR, evgA, kdpE, rstA* and *uvrY* fusions was not significantly changed, while that of *phoB* and *hprR* was slightly repressed, 1.6- and 1.4-fold, respectively. The strongest Hfq effect was observed on the *narP* fusion, since its activity was increased almost 3-fold in the *hfq* deletion strain. Interestingly, the expression of the *narP* RR gene was previously shown to be repressed at the post-transcriptional level by SdsN_137_, one isoform of a set of Hfq-dependent sRNAs involved in nitrogen metabolism [12]. However, SdsN sRNAs accumulate mostly in stationary phase as their transcription is dependent on the RpoS sigma factor and the level of SdsN_137_ is thus not expected to be maximum under the conditions used here. This makes it unlikely that the observed Hfq effect on *narP* is completely explained by the loss of regulation by SdsN_137_ in the *hfq* mutant (see also below), and suggests that other Hfq-dependent sRNAs might regulate *narP* expression.

### *narP* expression is modulated by several sRNAs

To identify these putative other sRNAs that might regulate *narP*, we made use of a plasmid library allowing overexpression of most Hfq-dependent sRNAs known to date. This library was initially created by Mandin & Gottesman [16], and was completed for the present study with plasmids overexpressing McaS [38], MicL [39], SdsN_137_ [12], CpxQ [40], DapZ [41], as well as the non Hfq-binding CsrB [42] sRNAs. The activity of the P_tet_-*narP-lacZ* fusion was thus measured in presence of all plasmids of the library (Fig. 2). The *narP* fusion is the same as that used in Table 1; it carries a 150 nt-long 5’ UTR that was chosen because this 5’ end was the most enriched after treatment with an exonuclease degrading 5’-monophosphate containing RNAs, and thus likely corresponds to the major *narP* TSS [37]. Four sRNAs had an effect greater than 2-fold: one positively (ChiX) and three negatively (DicF, RprA and SdsN_137_). Based on previous studies, the effect of ChiX is most probably due to a titration of Hfq [43,44]. For the repressing sRNAs, the effect of SdsN_137_ was expected and is in full agreement with the above-mentioned results [12]. DicF overproduction led to a strong growth defect and thus the possible repression of *narP* by DicF was not investigated further. The third repressing sRNA was RprA, whose transcription is under strong control by the Rcs phosphorelay [15]. RprA is known to activate the synthesis of the alternative sigma factor σ^S^ and of the RicI protein that inhibits conjugation [14,28]. Several negative targets of RprA have also been previously described: this sRNA represses expression of the *csgD* and *ydaM* (*dgcM*) genes involved in biofilm formation [27] and of *hdeD*, encoding an acid resistance protein [29]. Interestingly for this study, expression of other genes was found to be modulated in response to RprA pulse-overexpression in *Salmonella*, including *narP*, which was repressed [28].

**Figure 2:**
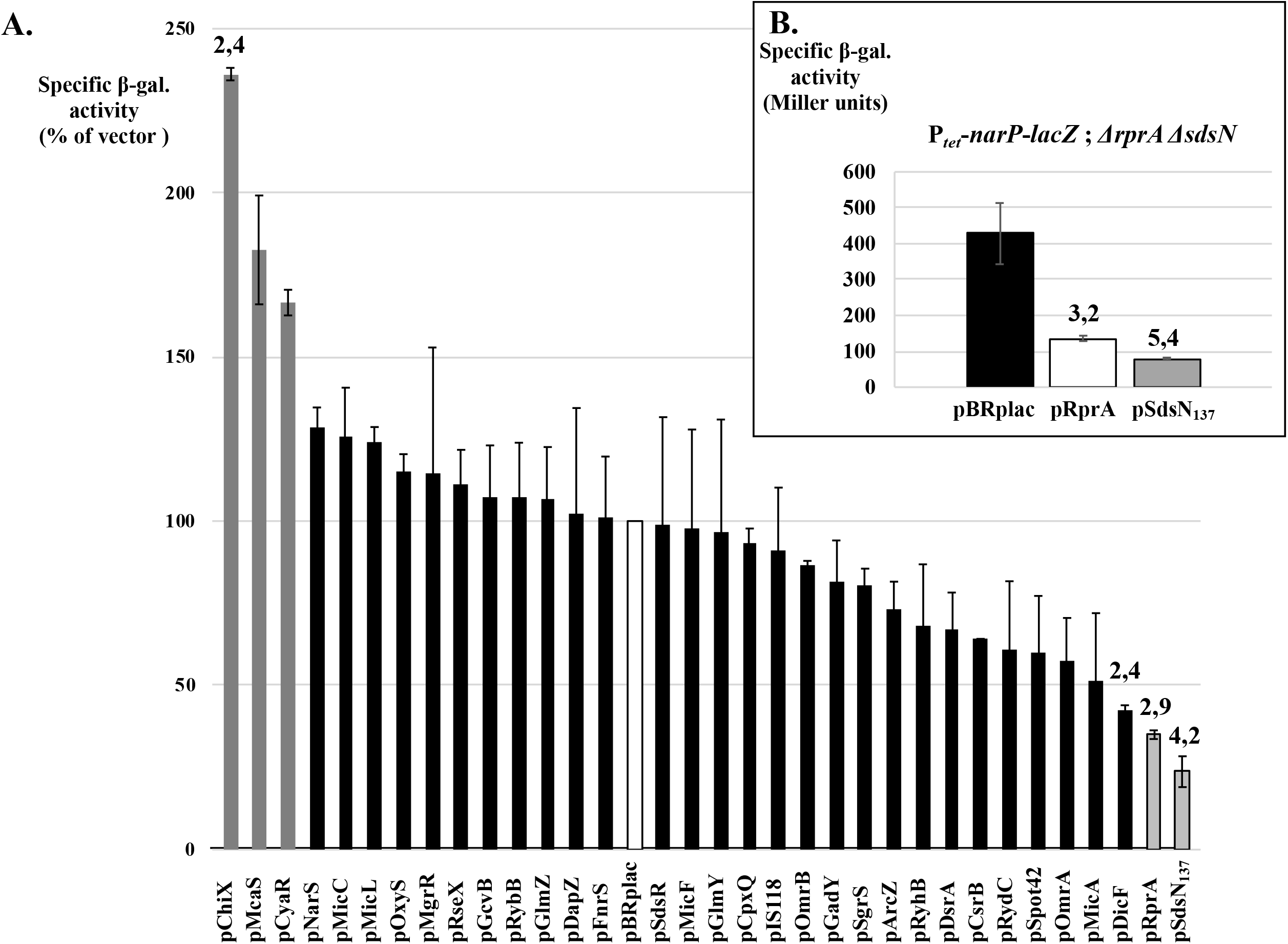
Hfq-dependent sRNAs modulate the expression of *narP*, most likely at the post-transcriptional level. (A) The β-galactosidase activity of the P_tet_-*narP-lacZ* fusion (strain AB1004) was measured in the presence of plasmids overexpressing most *E. coli* Hfq-dependent sRNAs reported to date. The activity of the fusion in presence of the vector control pBRplac was arbitrarily set at 100% and corresponds to an average value of 889 Miller units (with a standard deviation of 110) in LB-Amp_150_-IPTG. (B) The β-galactosidase activity of the same P_tet_-*narP-lacZ* fusion in a *ΔrprA ΔsdsN* background (strain AB1029) was measured in the presence of plasmids overproducing RprA or SdsN_137_ in LB-Amp_100_-IPTG.

Because RprA up-regulates RpoS synthesis, one might hypothesize that the pRprA plasmid used in Fig. 2A could promote SdsN_137_ synthesis by increasing σ^S^ levels, and thereby repress expression of the *narP-lacZ* fusion. However, the pRprA plasmid still regulated the *narP* fusion to a similar extent in a strain deleted for the chromosomal copy of *sdsN* and *rprA* (3.2-fold vs. 2.9-fold repression, Fig. 2B), indicating that its effect is independent of SdsN. We next investigated in more detail the control of *narP* by RprA.

### RprA directly targets the *narP* mRNA

Hfq-dependent sRNAs typically regulate gene expression by pairing to their targets, and we therefore looked for possible interactions between RprA and the *narP* mRNA using IntaRNA (Mann, Wright et al. 2017). The result is shown in Fig. 3A: nts 31-69 of RprA can potentially imperfectly base-pair to the TIR of *narP* messenger, from nts 116 to 154 relative to the *narP* TSS (i.e. nts -35 to +4 relative to *narP* start codon). Of note, affinity purification and sequencing of RNAs associated with a tagged version of RprA identified *narP* mRNA as a potential direct target, although not among the best candidates [29].

**Figure 3:**
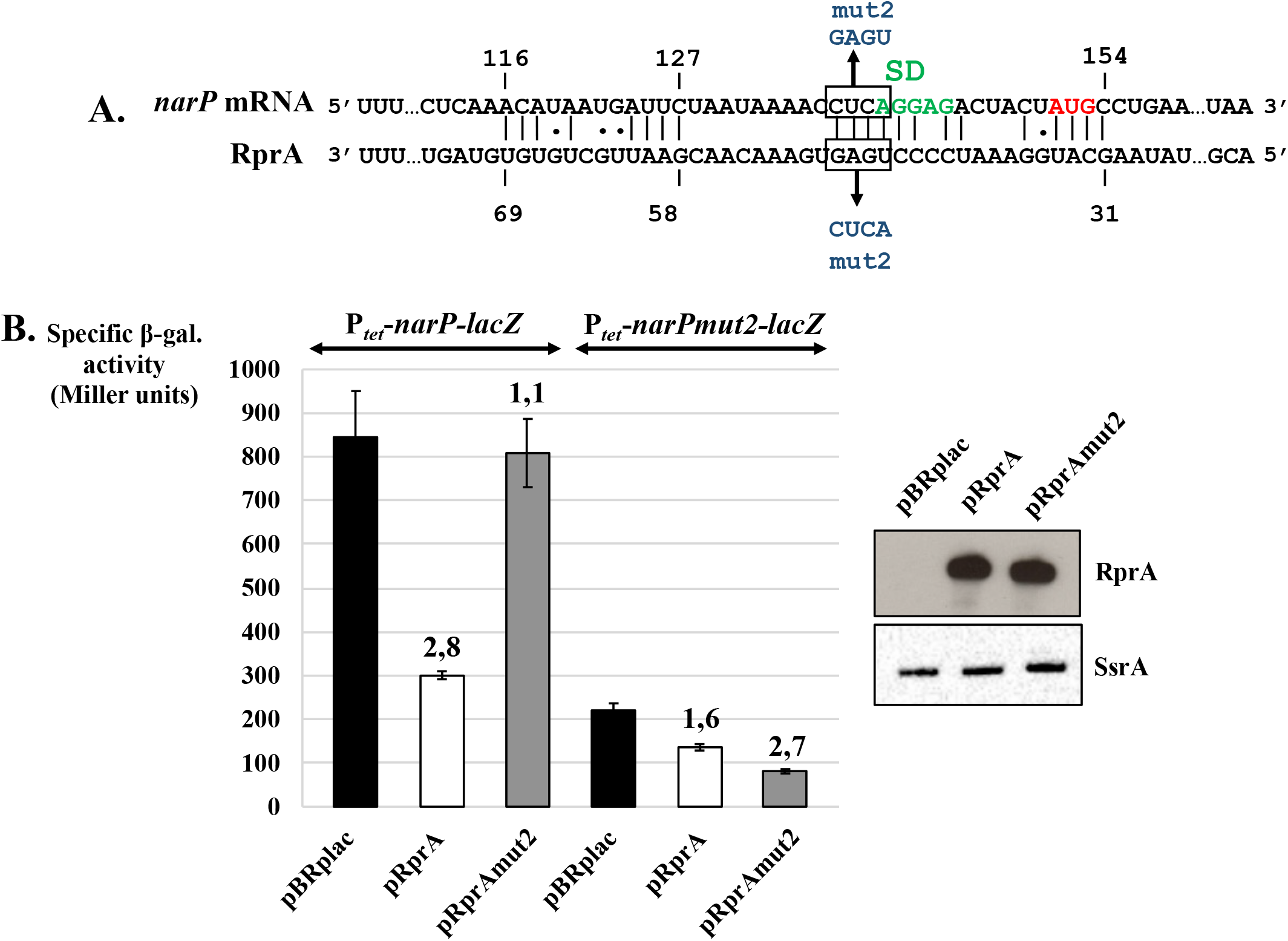
RprA directly pairs to the translation initiation region of *narP* mRNA. (A). Predicted base-pairing interaction between RprA and *narP* mRNA. The Shine-Dalgarno sequence and start codon of *narP* are indicated, and the compensatory mut2 changes in RprA or *narP-lacZ* fusion are shown in blue. (B) The β-galactosidase activity of the P_tet_-*narP*-*lacZ* translational fusion, wt or carrying the mut2 change, was measured upon overproduction of RprA or RprAmut2 in LB-Amp_150_-IPTG. Strains used in this experiment were AB1018 (wt fusion) and AB1037 (mut2) and are deleted for the chromosomal *rprA* copy. In parallel, RNA was extracted from strains with the wt fusion and Northern-blot analysis was done to assess the overproduction of RprA, wt or mut2, from pBRplac derivatives, using the RprA probe. SsrA was also probed from the same membrane and used as a loading control.

The RprA-*narP* mRNA predicted interaction (Fig. 3A) was experimentally tested by introducing mutations in RprA and/or in the previously described *narP-lacZ* fusion. Four possible bps adjacent or inherent to the Shine-Dalgarno (SD) sequence were disrupted by mutating either nts 45-48 of RprA from UGAG to ACUC (RprAmut2) or nts 137 to 140 of *narP-lacZ* from CUCA to GAGU (*narP*mut2). RprAmut2 abolished *narP* control, even though the sRNA accumulates to a level similar to that of the wt (Fig. 3B) while the mut2 change in *narP-lacZ* reduced control by RprA from 2.8-to 1.6-fold. A likely explanation for this residual control is proposed later in this study. Although the activity of the mutant fusion was strongly decreased, presumably because the mutation reduces the strength of the SD sequence, restoring the pairing by combining these compensatory changes restored control (2.7-fold repression, i.e. similar to the 2.8-fold repression of the wt fusion by wt pRprA), demonstrating that RprA sRNA directly pairs to *narP* mRNA *in vivo*. Since the validated interaction includes the Shine-Dalgarno sequence, RprA most likely inhibits *narP* translation initiation by preventing 30S ribosomal subunit binding to this mRNA.

### Overexpression of RprA or SdsN_137_ decreases NarP protein levels

In most cases, the active form of RRs is the phosphorylated form. However, it has been shown that robustness exists in TCS signaling and that changes in the total levels of a response regulator such as OmpR for instance, either by using an inducible heterogenous promoter or negative control by OmrA/B sRNAs, does not necessarily change the absolute levels of the phosphorylated form [17,45]. We thus wondered whether controlling *narP* synthesis by RprA and SdsN_137_ sRNAs would affect the levels of the phosphorylated form of the NarP RR (NarP-P). For this purpose, a tagged version of the NarP protein was constructed, where a 3xFlag sequence was added at its C-terminus after a short linker. This construction replaces the *narP* chromosomal copy. The biological activity of this tagged version of NarP was then assessed by measuring its ability to activate transcription from the *napF* promoter (Fig. 4A).

**Figure 4:**
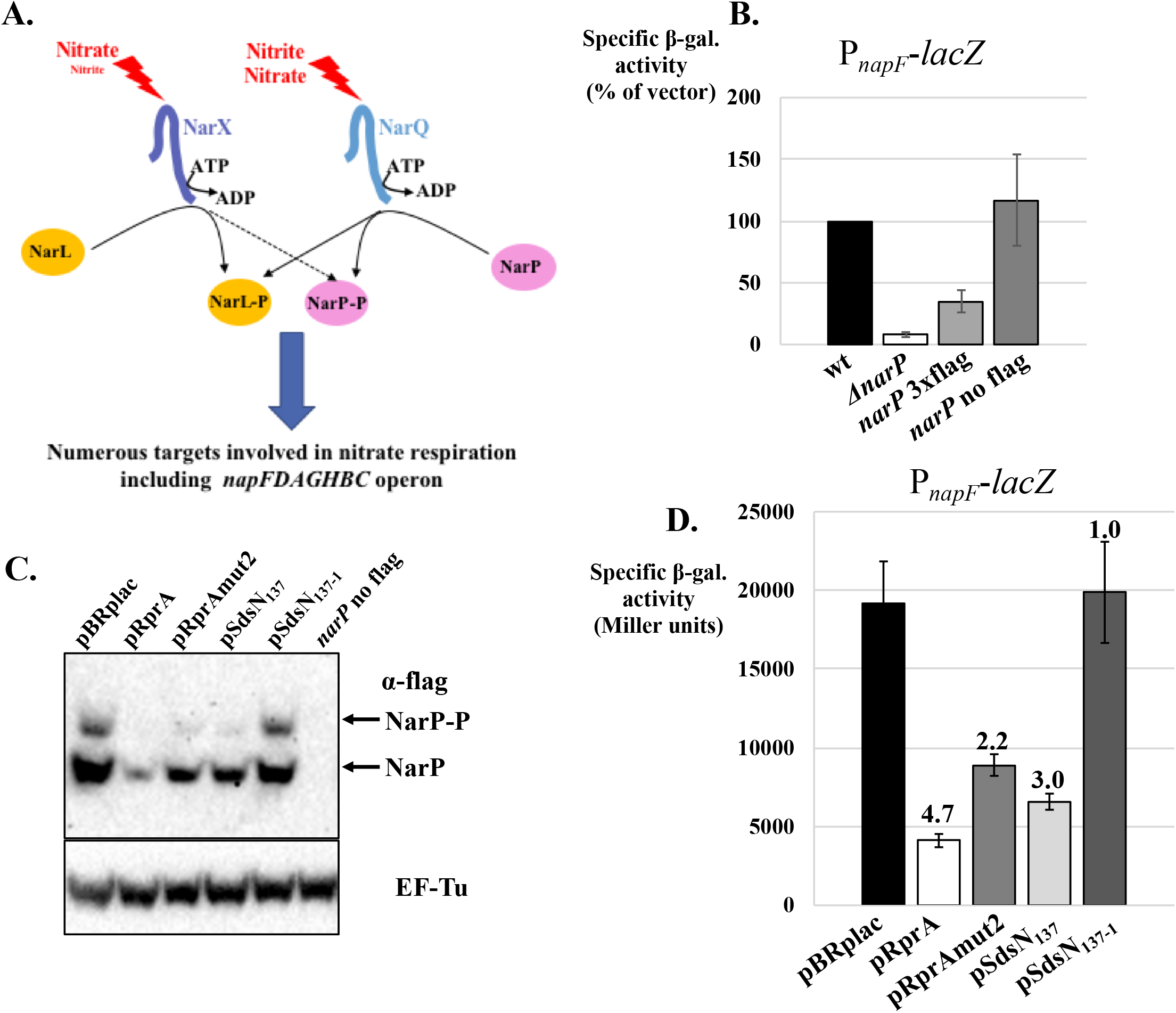
RprA and SdsN_137_ decrease the levels of both the phosphorylated and the non-phosphorylated forms of NarP and indirectly affect transcription of a NarP target. (A) Schematic of the regulatory network involving the two-component system NarQ-NarP. See text for details. (B) A NarP-3xFlag protein is partially active. Shown are the β-galactosidase activities of a transcriptional P_*napF*_-*lacZ* fusion in strains with a wt *narP* locus (strain AB1042), deleted of *narP* gene (AB1044), and where *narP* was replaced by a *narP*-linker-3xFlag construct followed by a kanamycin resistance gene (AB1082) or a control lacking the 3xFlag tag (no flag, strain AB1083). Activities were measured in MMGly+Ni under anaerobic condition. The activity of the fusion in the wt strain AB1042 was arbitrarily set at 100% and corresponds to 9774 Miller units in the first experiment and 21615 Miller units in the second. (C) Levels of the non-phosphorylated and phosphorylated forms of the NarP-3xFlag protein were followed by western-blot following protein separation on a Phos-Tag containing polyacrylamide gel. Protein samples were taken from cells grown in MMGly+Ni-Amp_100_-IPTG under anaerobic condition. Strains used for this experiment were AB1082, transformed with the indicated plasmids, or AB1083, transformed with the vector control, to ensure specificity of the Flag signal (see no Flag lane). EF-Tu levels were determined from the same membrane and used as a loading control. (D) β-galactosidase activity of the P_*napF*_*-lacZ* transcriptional fusion (in strain AB1042) was measured in the presence of plasmids overexpressing RprA and SdsN_137_, wt or mutated in the pairing region with *narP* translation initiation region. Cells were grown in MMGly+Ni-Amp_100_-IPTG under anaerobic condition.

The *E. coli napFDAGHBC* operon encodes the periplasmic nitrate reductase (and its accessory proteins) required for nitrate respiration in the presence of low concentration of this substrate [46]. Expression of the *napF* operon is induced by FNR and NarP-P, and repressed by NarL, as binding of NarL-P, the phosphorylated form of NarL to the *napF* promoter prevents binding of NarP-P. Because NarP and NarL are preferentially phosphorylated under low and high nitrate concentration, respectively, *napF* expression is expected to be higher under low nitrate conditions [47–49].

*napF* expression was thus followed by using a transcriptional fusion between the *napF* promoter (from nts -85 to +19 relative to the TSS from the proximal promoter) and *lacZ* sequence starting 17 nts before the translation initiation codon. In a preliminary experiment, the activity of this fusion was measured in cells grown under anaerobic conditions at different nitrate concentrations and was found to peak at around 5 mM nitrate (data not shown), in agreement with its control by NarL and NarP. The same minimal medium with glycerol as the sole carbon source and supplemented with 5mM nitrate (hereafter MMGly+Ni) was thus used in the next experiments. As expected, expression of the P_napF_*-lacZ* fusion was strongly decreased in the *narP* deleted strain (Fig. 4B). Furthermore, expression was partially restored in presence of the NarP-3xFLAG protein, showing that this tagged version of NarP retains biological activity, albeit at a reduced level compared to the wt protein (Fig. 4B).

The levels of NarP and NarP-P were then followed by western-blot using antibodies directed against the FLAG sequence upon overexpression of RprA and SdsN_137_ sRNAs, either wt or variants that are defective in *narP* control. The western-blots were performed by separating total proteins on a polyacrylamide gel containing Phos-Tag to allow the separation of NarP-P from the non-phosphorylated form of NarP and directly assess the effect of the sRNAs on the two forms of the protein. As shown in Fig. 4C, overexpression of wt RprA and SdsN_137_ significantly decreased the level of both forms of the NarP protein. Furthermore, this effect was abolished when we used instead the SdsN_137_-1 mutant that abolishes pairing to *narP* [12], indicating that these changes are due to *narP* post-transcriptional control by this sRNA. Strikingly however, RprAmut2, i.e. the RprA variant that no longer controls expression of the P_tet_-*narP-lacZ* fusion (Fig. 3B), was still very efficient at decreasing NarP and NarP-P levels (Fig. 4C and Fig. S2). This surprising effect of RprAmut2 is further discussed below (see Fig. 5). Changes in total NarP protein levels were also assessed in the same experiment by Western-Blot from a classical polyacrylamide gel where NarP and NarP-P are not separated; the results are in complete agreement with the Phos-Tag data and confirm the reduction in NarP levels in presence of pRprA, pRprAmut2, pSdsN_137_, but not pSdsN_137_-1 (Fig. S2).

**Figure 5.**
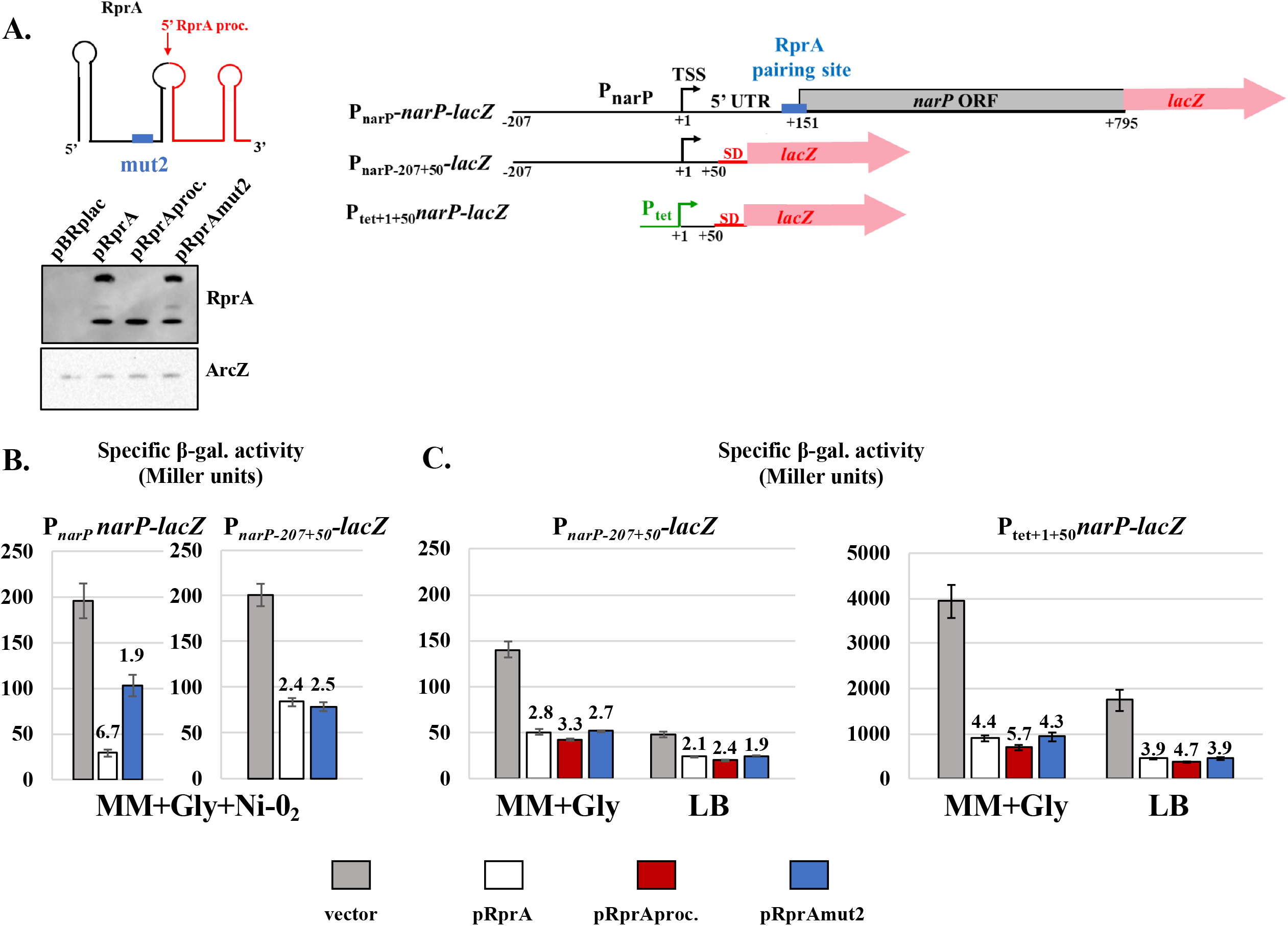
A second control of *narP* by RprA relies on the 5’ end of *narP* mRNA and the 3’ end of RprA. (A) Schematic of the different RprA versions and the three *narP-lacZ* fusions used in the next panels. (B, C) The β-galactosidase activity of the *narP-lacZ* fusions was measured upon overproduction of wt RprA or of several RprA variants, in MMGly-Amp_100_-IPTG medium (MM+Gly) or in LB-Amp-IPTG (LB). In panel B, cells were grown anaerobically in the presence of nitrate. Strains used in this experiment were AB1092 (P_narP_-*narP-lacZ* fusion), AB1159 (P_narP-207+50_-*lacZ*) and AB1184 (P_tet+1+50*narP*_-lacZ). In parallel to β-galactosidase assays of the right panel of Fig. 5C, total RNA was extracted from the same cultures in LB-Amp-IPTG, and RprA and ArcZ levels were analyzed by northern-blot using RprAproc and ArcZ biotinylated probes, respectively. The level of ArcZ was used as a loading control.

Overall, *narP* repression by RprA or SdsN_137_ results in a clear decrease of both NarP and NarP-P levels.

### Downstream effects of RprA and SdsN_137_ on NarP targets

Previous work has shown that regulating the synthesis of TCSs may have unexpected outcomes on the expression of their targets [45]. For instance, only one of the two sRNAs that repressed PhoP synthesis also repressed expression of PhoP-activated genes, while control of *ompR* by sRNAs affected only targets that were sensitive to the non-phosphorylated form of OmpR [17,18]. It was thus of interest to determine how RprA and SdsN_137_ control affected NarP-targets. For this purpose, we used the previously described P_napF_-*lacZ* transcriptional fusion and measured its activity upon overproduction of wt or mutant RprA and SdsN_137_ (Fig. 4D). Consistent with the observed reduction in NarP-P levels, RprA and SdsN_137_ decreased expression of the P_napF_ fusion by 4.7- and 3-fold, respectively. In contrast, SdsN_137_-1 had no effect, which was expected since this mutant no longer controls *narP*. The RprAmut2 variant still decreased expression of the P_napF_ fusion by 2.2-fold, which is fully consistent with its intermediary effect on the NarP protein levels (Fig. 4C). In other words, controlling NarP synthesis with RprA and SdsN_137_ sRNAs can impact NarP-targets, as shown here for *napF*. However, whether this is systematically true for other NarP targets remains to be determined.

### A second, independent, regulation of *narP* by RprA

The previous results clearly established the direct pairing between RprA and the *narP* translation initiation region (TIR) (Fig. 3). However, mutating RprA in this pairing region (RprAmut2) fully impaired control of a P_tet_-*narP-lacZ* fusion (Fig. 3B) but, surprisingly, did not completely abolish regulation of *narP* expression when looking at NarP protein levels and *napF* transcription (Fig. 4C and 4D). This could be explained if the mut2 change does not completely prevent the RprA interaction with *narP* TIR under some conditions, but we also considered the possibility that another regulation of *narP* by RprA may explain the residual control by RprAmut2. Because the latter effect was observed when *narP* was expressed from its own chromosomal locus in cells grown anaerobically in the presence of nitrate (Fig. 4), we first envisioned that this second control might target the *narP* promoter region and/or be observable only in specific experimental conditions.

To get further insight into this question, a *narP-lacZ* fusion where the P_tet_ promoter was replaced by the *narP* promoter was constructed (fusion P_narP_-*narP-lacZ* in Fig. 5A). Interestingly, the residual repression by RprAmut2 was also observed with this fusion, as RprAmut2 decreased its expression by almost 2-fold and wt RprA by more than 6-fold in MMGly+Ni under anaerobic conditions (Fig. 5B). We then used a fusion containing again the *narP* promoter but only the first 50 nts of the transcribed region of *narP* upstream of a *lacZ* gene carrying its own TIR (fusion P_narP-207+50_-*lacZ* in Fig. 5A). Even though the previously demonstrated interaction site with RprA is completely absent from this fusion, its expression was nonetheless found to be repressed to a similar extent by both wt RprA and RprAmut2 (2.4- and 2.5-fold repression, respectively, Fig. 5B). Thus, RprA also inhibits *narP* expression by acting either on its promoter or its early 5’ UTR, and this second control does not require the *narP* TIR. To determine whether this was observed only under anaerobic conditions in the presence of nitrate, the same experiment was repeated in aerobically-grown cells in MMGly and in LB medium (Fig. 5C, left panel). In both media, RprA and RprAmut2 repressed expression of the P_narP-207+50_-*lacZ* fusion to a similar extent: about 2.7-fold in MM+Gly and about 2-fold in LB. Hence, this second control of *narP* by RprA is not specific to anaerobic growth in the presence of nitrate. To explore this phenomenon further, we replaced the *narP* promoter region with a P_tet_ promoter (fusion P_tet+1+50narP_-*lacZ*, Fig. 5C, right panel). This did not alleviate control by RprA or RprAmut2, which clearly indicates that only the first 50 nts of the *narP* 5’ UTR are required for this second control by RprA. In addition, the overproduction of several other Hfq-binding sRNAs, including SdsN_137_, did not repress the expression of this P_tet+1+50narP_*-lacZ* fusion (data not shown), indicating that this effect is specific to RprA.

It is not clear at this stage why this second regulation did not allow control of the P_tet_-*narP-lacZ* fusion by RprAmut2 (Fig. 3B). It is possible that the combination of the experimental settings used in this experiment (aerobic growth in rich medium) with strong transcription from the P_tet_ promoter and the presence of the two regions of control, i.e. *narP* TIR and the first 50 nts of *narP* mRNA, is unfavorable for observing this regulation. Nevertheless, the residual control of the mutant *narPmut2-lacZ* fusion by the wt pRprA (Fig. 3B) is most likely explained by this second level of control acting on the (+1+50) region of *narP* mRNA.

Importantly, the effect of RprAmut2 and wt RprA on the shorter P_narP-207+50_- and P_tet+1+50narP_-*lacZ* fusions are similar. Thus, this second control most probably relies on a region of RprA that is not affected by the mut2 change, i.e. a different region than the one involved in the pairing to the *narP* TIR. Like many other sRNAs, RprA exists *in vivo* both as a full-length primary transcript, and as a shorter, processed form (RprA proc) corresponding to the last 47 nts of the sRNA [28,50]. Because RprAmut2 was as efficient as the wt in repressing expression of the fusions carrying only the first 50 nts of *narP* mRNA, we wondered whether this second control of *narP* could also be achieved by RprA proc. This would be consistent with the observation that RprAproc repressed *narP* in *Salmonella* even though this short version of the sRNA is devoid of the region interacting with *narP* TIR [28]. The activity of the fusions containing only the first 50 nts of *narP* mRNA was thus measured in presence of a plasmid overexpressing only the processed version of RprA (Fig. 5C): RprAproc was as efficient as wt RprA, and even more efficient in some cases, to control expression of these two fusions.

Together, these data clearly show that, in addition to the direct pairing of nts 30 to 70 of RprA to *narP* TIR, this sRNA also represses *narP* via a second action involving only the first 50 nts of *narP* mRNA and the 47 last nts of the sRNA.

### The Hfq effect on *narP-lacZ* fusion may be only partially due to sRNA regulation

As described above, we initially focused on the regulation of *narP* by sRNAs because *narP* expression was derepressed by about 3-fold in an *hfq*^*-*^ background (Fig. 1). Interestingly however, a similar increase in *narP* expression was observed in the *hfq* mutant in the absence of either *rprA* or *sdsN* genes, or in the double mutant, clearly showing that the Hfq effect is largely independent of these two sRNAs (Fig. 6A). This suggests that other Hfq-dependent sRNAs, not represented in the library used in Fig.2, could negatively regulate *narP* expression. Another possibility is that, in addition to being involved in RprA and SdsN action, Hfq could also directly modulate *narP* expression, this time independently of sRNAs, as shown for the *mutS* mRNA [7]. To discriminate between these two (non-exclusive) possibilities, we made use of point mutants of Hfq that have been previously described.

**Figure 6.**
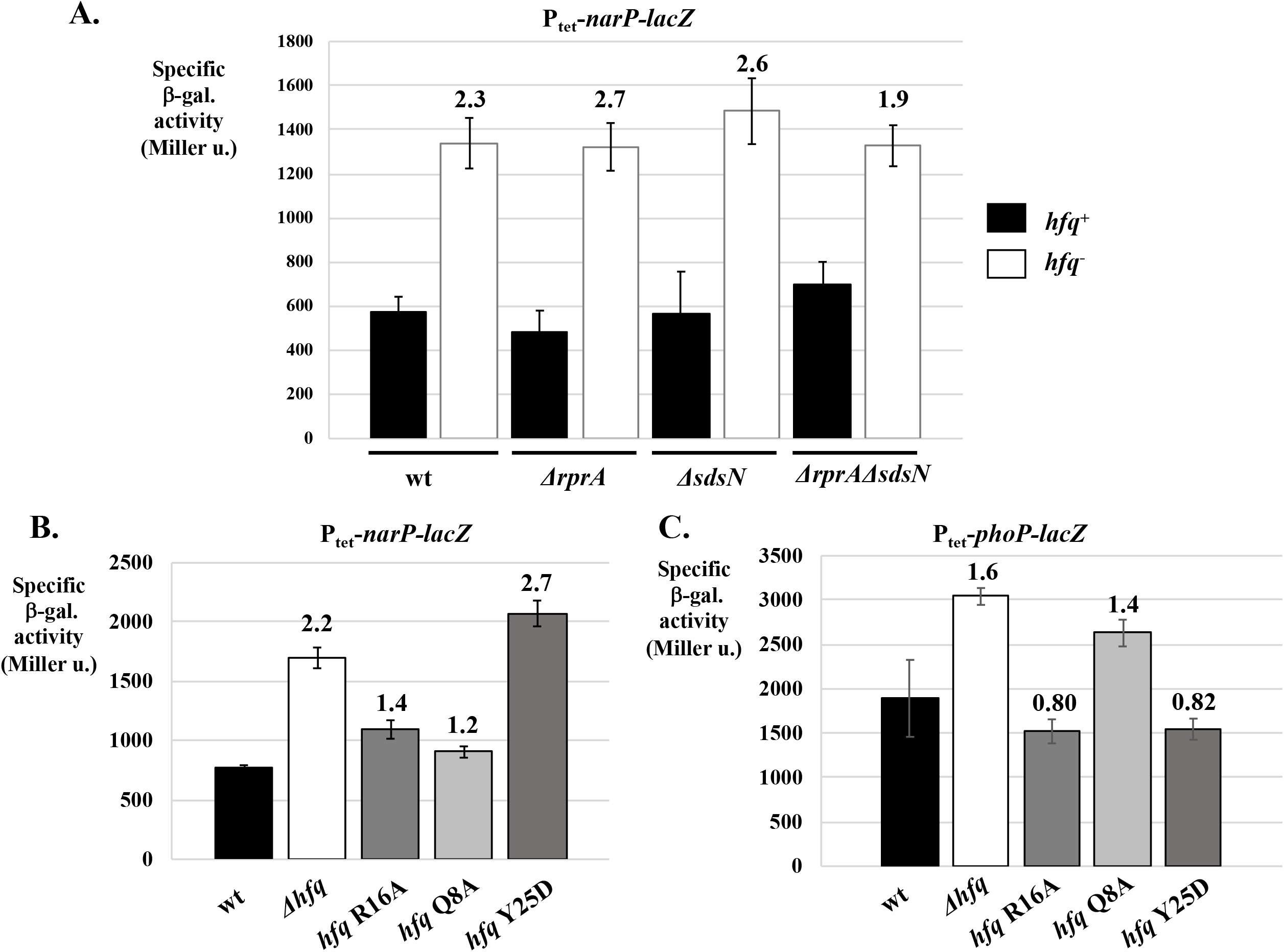
The Hfq effect on *narP* is largely independent of RprA and SdsN_137_ sRNAs. (A) The β-galactosidase activity of the P_tet_-*narP-lacZ* fusion was measured in LB medium in *hfq*^*+*^ and *hfq*^*-*^ background, in the presence and absence of the chromosomal copies of *rprA* and *sdsN* genes. Strains used here were AB1004, AB1013, AB1018, AB1109, AB1028, AB1110 AB1029 and AB1111. The effect of an *hfq* deletion or of mutations in the lateral face (R16A), proximal face (Q8A) or distal face (Y25D) of the Hfq protein was assessed on the activity of the same fusion (B) or that of the P_tet_-*phoP-lacZ* translational fusion (C). Strains used were AB1141, AB1142, AB1143, AB1144, AB1145, AB1151, AB1152, AB1153, AB1154 and AB1155.

Hfq forms a ring-shaped homohexamer that presents different surfaces involved in RNA binding: the proximal, distal, lateral faces and the C-terminal tail. In enterobacteria, several studies pointed to a major role of the proximal face in binding and stabilizing most sRNAs through interaction with the polyU stretch of the terminator, while the distal face displays a preference for (AAN) triplets, found in mRNAs and some sRNAs. The lateral face also participates to the binding of sRNAs or mRNAs via an arginine patch that can interact with UA-rich RNA sequences [31,51–55]. The expression of the P_tet_-*narP-lacZ* translational fusion was thus measured in point mutants affecting each of three surfaces: Q8A (proximal face), Y25D (distal face) and R16A (lateral face) [31]. Q8A and R16A mutants had no effect on the expression of the fusion, while the Y25D change caused a 2.7-fold increase, i.e. similar to the effect of the *hfq* null allele (Fig. 6B). These data suggest that, as for *mutS, narP* expression is not only subject to sRNA control, but is also likely directly controlled by Hfq. Since *phoP* expression was found to be up-regulated in the absence of Hfq, independently of the two known Hfq-dependent sRNAs that repress *phoP*, i.e. MicA and GcvB [18], the same set of mutants was tested with the P_tet_-*phoP-lacZ* fusion. In this case, the Q8A mimicked the effect of the *Δhfq* mutation, while R16A and Y25D had no effect, consistent with the hypothesis that *phoP* expression is regulated by Hfq-dependent sRNAs that remain to be identified.

## DISCUSSION

### A new connection between sRNAs and nitrate metabolism

Enterobacteria such as *Escherichia coli* and *Salmonella* are facultative anaerobes and often encounter low oxygen conditions, e.g. in the gut of mammalian hosts. Their ability to respire on nitrate or nitrite, which are efficient alternative electron acceptors to oxygen, is certainly an advantage under anaerobic conditions and this has been linked to efficient colonization [56] and competitive growth in mice large intestine during the host inflammatory response, which generates nitrate [57]. Several transcriptional regulators ensure control of gene expression in response to the availability of different electron acceptors. In particular, FNR and ArcB-ArcA both respond to anaerobiosis, with the oxidation status of quinones being involved in signaling to ArcB-ArcA. The NarX-NarL and NarQ-NarP TCSs are involved in the response to nitrate and nitrite. While the NarQ sensor phosphorylates both the NarP and NarL RR in response to nitrate and nitrite, NarX preferentially phosphorylates NarL in response to nitrate mostly [58,59]. NarL and NarP regulons partially overlap and include genes for enzymes required for nitrate/nitrite respiration, among them the nitrate reductases NarGHJI (membrane-bound) and NapFDAGHBC (periplasmic), and the nitrite reductases NirBDC and NrfABCDEFG. Other members of the NarP and the NarL regulons, e.g. the *hcp-hcr* operon, are involved in the response to nitrogen stress and NO detoxification following nitrate respiration [60–63]. The identification of *narP* as one of the direct targets of RprA suggests that it may be advantageous for the cell to limit NarP levels under conditions where RprA is expressed, such as cell surface stress that would signal to the Rcs phosphorelay [64]. In line with this, transcriptomic analyses of strains lacking the RcsB regulator identified several genes related to nitrogen metabolism, e.g. *napAB, nirBDC, narK* or *narGH*, as repressed by the Rcs system in several bacteria [33,65].

Reducing NarP levels could facilitate the activation of NarL by NarX or NarQ; this would be true for control by either RprA or SdsN_137_. Such a precise balance between NarP and NarL functions may be important for a proper response to different nitrate concentrations; for example, NarX-NarL has been found to be more important for intestine colonization than NarQ-NarP, possibly because it corresponds to relatively high nitrate conditions where the NarGHI reductase, whose expression is activated by NarL, plays a key role [56]. In this regard, it is interesting to note that SdsN_137_ levels vary in response to some nitrogen sources [12]. Even though RprA and SdsN_137_ production may not peak under the same conditions, the two sRNAs can be co-expressed, for example in stationary phase where they are both detected [12,50,66], which could contribute to stronger reduction of *narP* expression under such conditions.

Interestingly for this study, *narL* could also be targeted by sRNAs as its mRNA levels were shown to be decreased upon overexpression of DicF, an sRNA that accumulates under micro-aerobic conditions [67], and that was identified here as a possible regulator of *narP* as well (Fig. 2). Furthermore, NarL also promotes synthesis of the NarS sRNA, which is processed from the NarL-activated *narK* mRNA, encoding a nitrate/nitrite antiporter [68]. In turn, NarS represses the expression of the gene encoding the nitrite transporter NirC. Although this NarS-control does not affect the expression of the other genes of the *nirBDC-cysG* operon, the synthesis of the NirB subunit of the NADH-dependent nitrite reductase is also subject to sRNA control, in this case by the RyhB sRNA [69]. The link between sRNAs and nitrogen metabolism is further illustrated by the fact that SdsN_137_ represses the synthesis of the NfsA nitroreductase and the HmpA nitric oxide dioxygenase [12]. Additionally, several global approaches looking at ProQ or Hfq targets, or at RNA-RNA interactions mediated by these chaperones, indicate that yet other genes related to nitrate/nitrite metabolism are likely to be controlled by sRNAs in enterobacteria [30,34,35].

Lastly, connections between sRNAs and nitrogen are not restricted to enterobacteria and many other examples have been reported in extremely diverse bacteria as well, including the sRNAs related to denitrification in *Pseudomonas aeruginosa* or *Paracoccus denitrificans* [70,71], the RoxS sRNA that responds to nitric oxide in firmicutes [72] and the sRNAs involved in the control of carbon-nitrogen balance in cyanobacteria [73].

### New connections between Hfq and two-component systems

This study adds RprA to the list of sRNAs that include genes for TCSs in their regulons. Previous work has shown that altering the levels of transcriptional regulators with sRNAs may not always lead to a change in their activity, especially in the case of TCS regulators that must be activated by phosphorylation. In particular, the EnvZ-OmpR TCS was found to be robust, i.e. the level of OmpR-P, the phosphorylated form of OmpR, is insensitive to large changes in total EnvZ or OmpR levels [45]. Consistent with this, repressing *ompR* expression with OmrA and OmrB sRNAs decreased only the amount of the non-phosphorylated form of OmpR, which allowed these sRNAs to indirectly limit their own synthesis as their transcription responds to both the phosphorylated and the non-phosphorylated forms of OmpR [17]. Although different, the outcomes of controlling PhoP synthesis with MicA or GcvB sRNA were also surprising: of these two sRNAs that repressed *phoP* via competition with ribosome binding, only MicA decreased the expression of positive PhoP-targets [18].

In the case of *narP*, our results show that repression by RprA or SdsN_137_ sRNAs decreased the level of both the phosphorylated and the non-phosphorylated form of this RR and, consistently, repressed transcription from the NarP-dependent promoter P_napF_. This sRNA effect on the phosphorylated form of the RR differs from what has been observed for *ompR* and indicates that robustness is not true for all TCS. However, for most of the previously reported cases, genes for RRs whose expression is repressed by sRNAs are in an operon with their cognate sensor kinase genes, leading to the prediction that expression of the kinase would also be repressed by the sRNA. This is different for *narP* since this is one of the few *E. coli* RR genes that is not part of an operon. Because RprA does not appear to control *narQ* (our preliminary data), its control of *narP* should thus change the RR/SK ratio. It would be interesting to determine whether this explains the observed difference in the levels of phosphorylated forms of NarP and OmpR in response to repression by sRNAs, which are decreased or unaffected, respectively (this study and [17]).

Another conclusion of this study is that, in addition to allowing sRNAs function, Hfq could also play a direct role in controlling TCSs expression. Although more direct experiments are required to definitively show that the effect of the Hfq Y25D mutant on *narP* is due to a defect in mRNA binding, and that sRNAs are not involved, this nonetheless indicates that *narP* expression could vary in response to signals that affect Hfq synthesis, stability and/or activity. While a similar direct Hfq effect on *phoP* seems unlikely at this stage, these data also raise the question of whether other TCSs can be controlled by Hfq independently of sRNAs, as previously shown for *mutS* [7].

In general, it will be important to determine how these post-transcriptional control mechanisms involving Hfq and/or sRNAs affect the various TCS signaling properties and, especially in the case of NarQ-NarP, the crosstalk with other systems.

### Two independent actions of RprA to control narP

This study is also interesting from a mechanistic standpoint, with the finding that a sRNA can repress a single target via two pathways, involving different regions of both the sRNA and the mRNA. The first, canonical, control relies on the pairing of RprA to the *narP* TIR, which presumably blocks translation initiation. The second control only requires the first 50 nts of *narP* mRNA, i.e. a region that is distant from the TIR since *narP* 5’UTR is 150 nt-long. These two pathways are most likely independent as the 5’end region of *narP* mRNA is sufficient to observe regulation by RprA in the absence of the TIR (Fig. 5) and, similarly, RprA efficiently represses expression of the P_BAD_-*narP-lacZ* fusion used in [12] that carries a 78-nt 5’ UTR and is thus devoid of the 5’ end of the *narP* 5’ UTR used here (our unpublished results).

The second control can be performed with similar efficiencies by either the full-length or the processed form of RprA carrying only the last 47 nts of the sRNA. Because we could not predict a convincing base-pairing between the 5’end of *narP* messenger (*narP*(+1+50) region) and RprA proc, this second control is likely not due to a direct sRNA-mRNA interaction. Instead, one can envision that RprA, or RprA proc, would control the level or the activity of a factor that would act on the *narP*(+1+50) region. A first possibility is that this factor mediating the observed control of RprA on *narP*(+1+50) is another sRNA. Such a scenario would be similar to other regulatory circuits involving sponge RNAs (here, RprA) that can titrate and/or destabilize an sRNA and thereby prevent its action [74,75]. We have already tested a possible role of a few candidates sRNAs in the *narP* control by RprA. These candidates were sRNAs whose levels vary according to aerobic/anaerobic conditions (FnrS, ArcZ, or NarS), sRNAs predicted to interact with RprA based on RIL-seq data (sRNA from the *ariR-ymgC* intergenic region), or sRNAs whose expression could be related to that of *narP* (narP 3’UTR). However, our results so far do not support the involvement of any of these sRNAs in this control (our unpublished data). Although possible factor(s) intervening in the effect of RprA on *narP*(+1+50) remain to be precisely identified, the dual-control mechanism analyzed here is most likely different from the reported examples of RprA pairing to two distinct sites on a single target, such as *csgD* [27] or *hdeD* [29].

Another question raised by our results is that of the step at which this second control takes place. Given that the promoter and the TIR of *narP* are dispensable for this regulation, it is very unlikely to occur at the level of initiation of *narP* transcription or translation. However, an 8-aa upstream ORF is predicted in the first 50 nts of *narP* and its translation could be the target of this second control. Alternatively, the action on *narP*(+1+50) could primarily affect the stability of *narP* mRNA, as previously reported for other bacterial sRNAs, possibly by targeting the very 5’ end of mRNAs [76– 78]. The effect could also be transcriptional, for instance if RprA indirectly induced a premature termination of *narP* transcription.

There are several examples of feed-forward regulatory motifs where sRNAs regulate the expression of a target both directly and indirectly; the indirect effect is often mediated by a transcriptional regulator acting on the target promoter. One such example is the activation of the synthesis of the conjugation-inhibiting protein RicI by RprA that relies both on the pairing of RprA to *ricI* mRNA and on RprA promoting translation of the RpoS sigma factor that activates *ricI* transcription [28]. It is likely that the dual control of *narP* by RprA reported here will also result in a feed-forward regulatory circuit, although with different details. Identifying the missing clues of this circuit, and in particular the nature of the indirect factor(s) involved and the way it acts, will be crucial to assess how this dual control can precisely impact *narP* expression.

Regardless of the precise mechanism of this second control, it is clear that the processed version of the sRNA is sufficient to promote it, even though this does not necessarily tell us whether the full-length or the processed form of RprA is preferentially responsible for this regulation when they are both present in the cell. RprA is not the only sRNA that exists under different isoforms, either because of processing, leading for instance to short forms of ArcZ, SdsR, MicL, RbsZ just to name a few [16,35,39,79,80], or of multiple TSS, which explains the different SdsN species [12]. In the future, it will be interesting to decipher whether full-length and short forms of other sRNAs can also complement each other in the regulation of some targets, as reported here for RprA and *narP*.

## ACKNOWLEDGMENTS

We thank G. Storz, N. Majdalani, S. Gottesman, P. Mandin, E. Solchaga and J. Plumbridge for providing many strains and plasmids in the course of this study. We are grateful to P. Mandin (LCB) for help with RNA extraction from anaerobically-grown cells and to members of the Magalon’s lab, especially L. Pieulle, F. Seduk and S. Bulot for assistance. We thank members of the group at the IBPC for support and discussions, and M. Springer, S. Gottesman, C. Condon and C. Chiaruttini for insightful comments on the manuscript.

---

**Figure S1: Overproduction of DapZ and CpxQ**

The levels of DapZ (A) and CpxQ (B) sRNAs were assessed by Northern-blot after RNA extraction from NEB5α-F’I^q^ strain transformed by pBRplac empty vector, or its derivatives carrying either sRNA gene. Cells were grown in LB Amp IPTG 10^−4^M. The level of SsrA was also analyzed and used as a loading control.

**Figure S2: RprA and SdsN**_**137**_ **decrease the levels of NarP**

Levels of NarP-3xFlag protein were analyzed by western-blot following protein separation on a standard polyacrylamide gel (A) or on Phos-Tag containing gels (B & C). Samples used in the western-blot of panel (A) are the same as in figure 4C. Panel (B) is an independent biological replicate from the experiment presented in Fig. 4C. Panel (C) displays the entire gel shown in Fig. 4C. Asterisks indicate samples that have been diluted 2- or 4-fold from the “pBRplac” sample of the first lane; “+0_2_” corresponds to a sample from cells transformed with the pBRplac empty vector and grown under aerobic conditions.

