## Supplementary figures and images for "Synthesis of the NarP response regulator of nitrate respiration in *Escherichia coli* is regulated at multiple levels by Hfq and small RNAs"

### Fig. S1

#### Slide 1
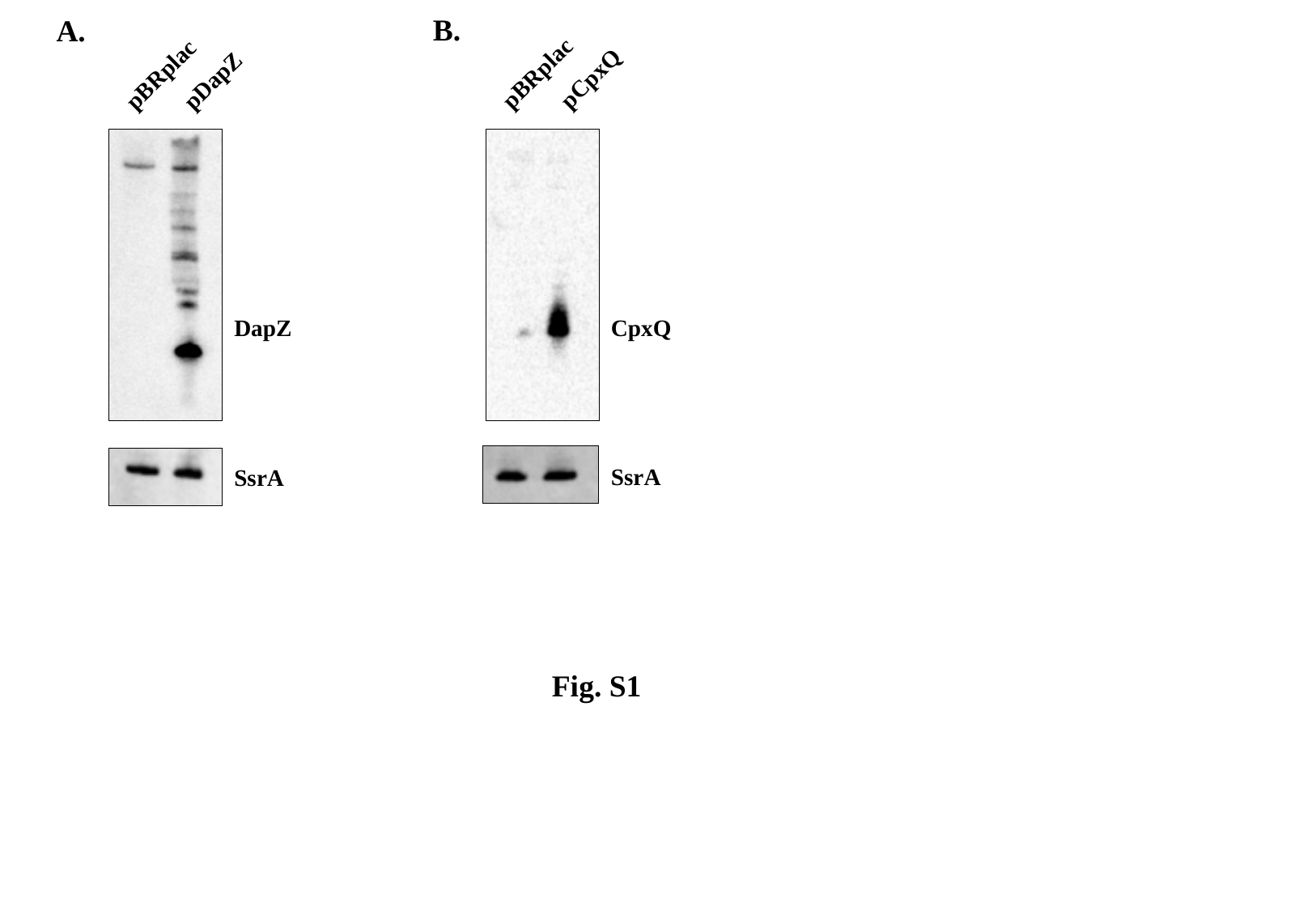

B.
A.
pCpxQ
pDapZ
pBRplac
pBRplac
DapZ
CpxQ
SsrA
SsrA
Fig. S1
