## Supplementary material for "Synthesis of the NarP response regulator of nitrate respiration in *Escherichia coli* is regulated at multiple levels by Hfq and small RNAs": Fig. S2

### Slide 1
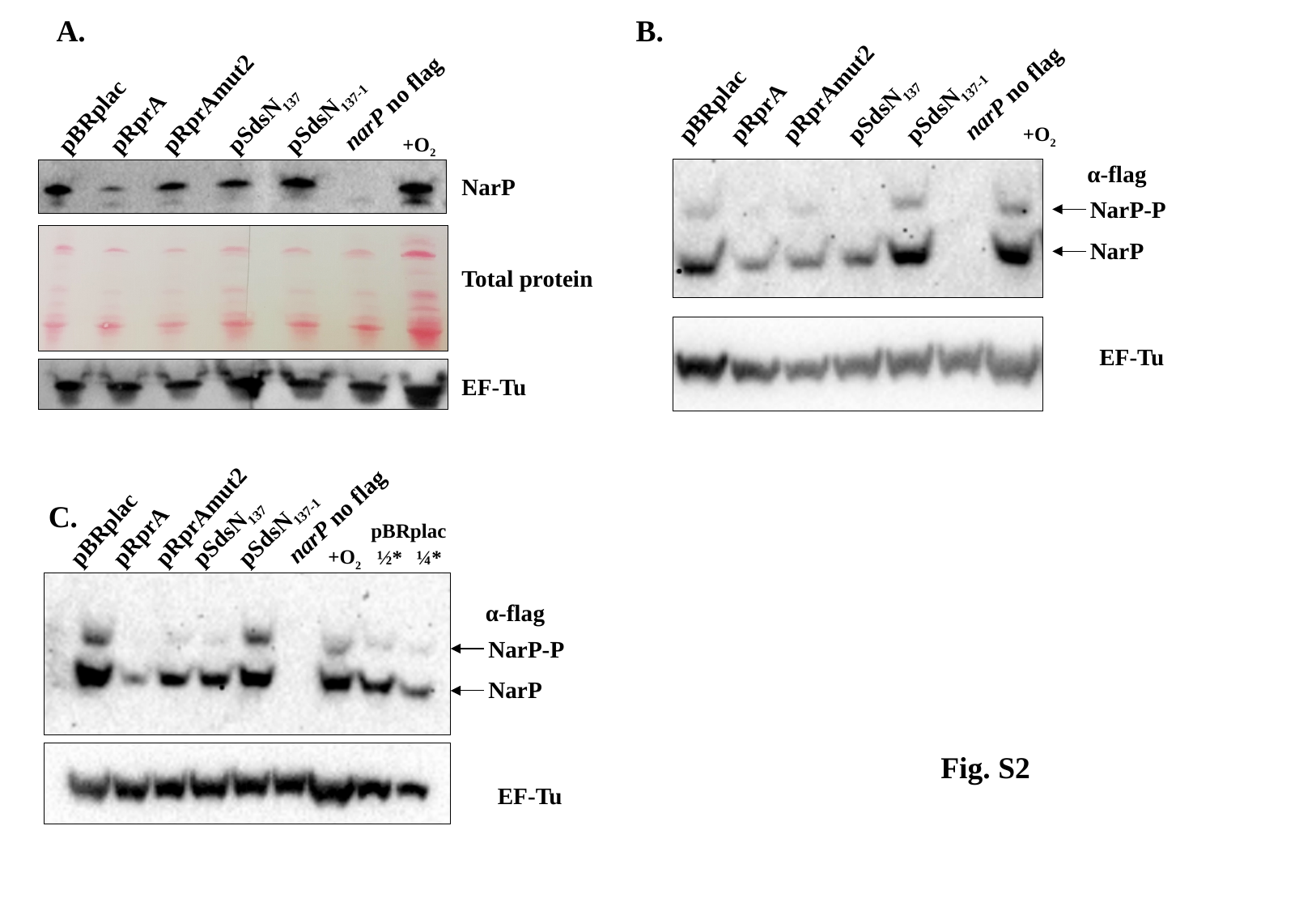

A.
B.
pRprAmut2
narP no flag
pSdsN_137-1_
pSdsN_137_
pBRplac
pRprA
+O_2_
NarP
Total protein
EF-Tu
narP no flag
pRprAmut2
pSdsN_137-1_
pSdsN_137_
pBRplac
pRprA
+O_2_
α-flag
NarP-P
NarP
EF-Tu
pRprAmut2
narP no flag
pSdsN_137-1_
pSdsN_137_
pBRplac
pBRplac
pRprA
+O_2_
½*
¼*
α-flag
NarP-P
NarP
EF-Tu
C.
Fig. S2
