## Supplementary material for "Synthesis of the NarP response regulator of nitrate respiration in *Escherichia coli* is regulated at multiple levels by Hfq and small RNAs": Table S1

### Table S1: Strains and plasmids used in this study

| Strain name | Characteristics | References |
|---|---|---|
| MG1655 | *E.coli* K-12 wt strain | Strain obtained from F. Blattner's lab |
| DJ624 | MG1655 *ΔlacX74 mal ::lacIq* | D. Jin, NIH |
| MG1110 | DJ624 *hfq ::cat* | P1, (*hfq ::cat* mutant from (1)) into DJ624 |
| MG1433 | DJ624 *mini-λ-cmp*$^R$ | DJ624 + P1 (NM1100, from N. Majadalani, unpublished) |
| GSO762 | MG1655 Δ*sdsN::kan* | (2) |
| JW2181 | *narP ::kan* mutant from the Keio collection | (3) |
| DJS2604 | MG1655 *Δhfq ::cat-sacB purA ::kan* | From D. Schu, unpublished |
| DJS2609 | MG1655 *Δhfq purA*$^+$ | From D. Schu, unpublished |
| DJS2927 | MG1655 *hfq*$^+$ *purA*$^+$ | From D. Schu, unpublished |
| KK2560 | MG1655 *hfqQ8A purA*$^+$ | From D. Schu, unpublished |
| KK2561 | MG1655 *hfqR16A purA*$^+$ | From D. Schu, unpublished |
| KK2562 | MG1655 *hfqY25D purA*$^+$ | From D. Schu, unpublished |
| MG1508 | MG1655 *mal ::lacI*$^q$ *mini-λ-tet*$^R$ $P_{tet}$-*cat-sacB-lacZ*$_{+28}$ | (4) |
| MG1511 | MG1508 $P_{tet}$-*phoP*$_{-36+30}$-*lacZ*$_{+28}$ | (4) |
| MG1515 | MG1511 *hfq ::cat* | (4) |
| MG2114 | MG1655 *mal ::lacI*$^q$ *mini-λ-tet*$^R$ *rrnBt2-cat-sacB-lacZ*$_{-17}$ | This study |
| AB1000 | MG1508 $P_{tet}$-*baeR*$_{-30+720}$-*lacZ*$_{+28}$ (numbering relative to start codons) | This study, Recombineering in MG1508 |
| AB1001 | MG1508 $P_{tet}$-*basR*$_{-18+666}$-*lacZ*$_{+28}$ (numbering relative to start codons) | This study, Recombineering in MG1508 |
| AB1002 | MG1508 $P_{tet}$-*evgA*$_{-125+612}$-*lacZ*$_{+28}$ (numbering relative to start codons) | This study, Recombineering in MG1508 |
| AB1003 | MG1508 $P_{tet}$-*kdpE*$_{-39+675}$-*lacZ*$_{+28}$ (numbering relative to start codons) | This study, Recombineering in MG1508 |
| AB1004 | MG1508 $P_{tet}$-*narP*$_{-150+645}$-*lacZ*$_{+28}$ (numbering relative to start codons) | This study, Recombineering in MG1508 |
| AB1005 | MG1508 $P_{tet}$-*phoB*$_{-41+687}$-*lacZ*$_{+28}$ (numbering relative to start codons) | This study, Recombineering in MG1508 |
| AB1006 | MG1508 $P_{tet}$-*rstA*$_{-13+726}$-*lacZ*$_{+28}$ (numbering relative to start codons) | This study, Recombineering in MG1508 |
| AB1007 | MG1508 $P_{tet}$-*uvrY*$_{-44+654}$-*lacZ*$_{+28}$ (numbering relative to start codons) | This study, Recombineering in MG1508 |
| AB1008 | MG1508 $P_{tet}$-*hprR*$_{+1+669}$-*lacZ*$_{+28}$ (numbering relative to start codons) | This study, Recombineering in MG1508 |
| AB1009 | AB1000 *hfq ::cat* | This study, AB1000 + P1 (MG1110) |
| AB1010 | AB1001 *hfq ::cat* | This study, AB1001 + P1 (MG1110) |
| AB1011 | AB1002 *hfq ::cat* | This study, AB1002 + P1 (MG1110) |
| AB1012 | AB1003 *hfq ::cat* | This study, AB1003 + P1 (MG1110) |
| AB1013 | AB1004 *hfq ::cat* | This study, AB1004 + P1 (MG1110) |
| AB1014 | AB1005 *hfq ::cat* | This study, AB1005 + P1 (MG1110) |
| AB1015 | AB1006 *hfq ::cat* | This study, AB1006 + P1 (MG1110) |
| AB1016 | AB1007 *hfq ::cat* | This study, AB1007 + P1 (MG1110) |
| AB1017 | AB1008 *hfq ::cat* | This study, AB1008 + P1 (MG1110) |
| AB1018 | AB1004 *rprA ::tet* | AB1004+P1(NM667, from N. Majdalani, unpublished) |
| AB1028 | AB1004 *sdsN ::kan* | This study, AB1004+P1(GS0762) |
| AB1029 | AB1018 *sdsN ::kan* | This study, AB1018+P1(GS0762) |
| AB1037 | MG1508 $P_{tet}$-*narP*$_{-150+645}$<u>*mut2*</u>-*lacZ*$_{+28}$ *rprA ::tet* (numbering relative to start codons) | This study, Recombineering in MG1508 |
| AB1042 | MG2114 *rrnBt2*-$P_{napF-85+19}$-*lacZ*$_{-17}$ (numbering of *napF* relative to its TSS and of *lacZ* relative to its start codon) | This study, Recombineering in MG2114 |
| AB1044 | AB1042 *narP ::kan* | This study, AB1042+P1(JW2181) |
| AB1070 | MG1433 *narP-linker-3xflag-kanR* | This study, Recombineering in MG1433 |
| AB1073 | MG1433 *narP-linker-kanR* | This study, Recombineering in MG1433 |
| AB1082 | AB1042 *narP-linker-3xflag-kanR* | This study, AB1042+P1(AB1070) |
| AB1083 | AB1042 *narP-linker-kanR* | This study, AB1042+P1(AB1073) |
| AB1092 | MG1508 *rrnBt2*-$P_{narP}$-*narP*$_{-207+795}$-*lacZ*$_{+28}$ (numbering of *narP* relative to its TSS and of *lacZ* relative to its start codon) | This study, Recombineering in MG1508 |
| AB1109 | AB1018 *hfq ::cat* | This study, AB1018+P1(MG1110) |
| AB1110 | AB1028 *hfq ::cat* | This study, AB1028+P1(MG1110) |
| AB1111 | AB1029 *hfq ::cat* | This study, AB1029+P1(MG1110) |
| AB1127 | AB1004 *Δhfq :: cat-sacB purA :: kan* | This study, AB1004+P1(DJS2604) |
| AB1141 | AB1127 *hfq*$^+$ *purA*$^+$ | This study, AB1127+P1(DJS2927) |
| AB1142 | AB1127 *Δhfq purA*$^+$ | This study, AB1127+P1(DJS2609) |

| | | |
|---|---|---|
| AB1143 | AB1127 *hfq*R16A *purA*$^{+}$ | This study, AB1127+P1(KK2561) |
| AB1144 | AB1127 *hfq*Y25D *purA*$^{+}$ | This study, AB1127+P1(KK2562) |
| AB1145 | AB1127 *hfq*Q8A *purA*$^{+}$ | This study, AB1127+P1(KK2560) |
| AB1148 | MG1508 Ptet-*ompR*$_{-35+30}$-*lacZ*$_{+28}$ (numbering relative to start codons) | This study, Recombineering in MG1508 |
| AB1149 | AB1148 *hfq ::cat* | This study, AB1148+P1(MG1110) |
| AB1150 | MG1511 *Δhfq* :: *cat-sacB purA :: kan* | This study, MG1511+P1(DJS2604) |
| AB1151 | AB1150 *hfq*$^{+}$ *purA*$^{+}$ | This study, AB1150+P1(DJS2927) |
| AB1152 | AB1150 *Δhfq purA*$^{+}$ | This study, AB1150+P1(DJS2609) |
| AB1153 | AB1150 *hfq*R16A *purA*$^{+}$ | This study, AB1150+P1(KK2561) |
| AB1154 | AB1150 *hfq*Y25D *purA*$^{+}$ | This study, AB1150+P1(KK2562) |
| AB1155 | AB1150 *hfq*Q8A *purA*$^{+}$ | This study, AB1150+P1(KK2560) |
| AB1159 | MG1508 *rrnBt2*-P$_{narP-207+50}$-*lacZ*$_{-17}$ (numbering of *narP* relative to its TSS and of *lacZ* relative to its start codon) | This study, Recombineering in MG1508 |
| AB1184 | MG1508 *rrnBt2*-Ptet$_{+1+50}$*narP*-*lacZ*$_{-17}$ (numbering of *narP* relative to its TSS and of *lacZ* relative to its start codon) | This study, Recombineering in MG1508 |
| **Plasmid** | **Characteristics** | **Use** |
| pBRplac | Ampicillin$^{R}$, tetracycline$^{R}$, P$_{LlacO-1}$ cloned into pBR322 | Vector control, from (5) |
| pRprA | *rprA* gene under P$_{LlacO-1}$ control in pBRplac | RprA overproduction, from (6) |
| pRprAmut2 | *rprAmut2* under P$_{LlacO-1}$ control in pBRplac | RprAmut2 overproduction, this study |
| pRprAproc | *rprAproc* under P$_{LlacO-1}$ control in pBRplac | RprAproc overproduction, this study |
| pSdsN$_{137}$ | *sdsN*$_{137}$ under P$_{LlacO-1}$ control in pBRplac | SdsN$_{137}$ overproduction, from (2) |
| pSdsN$_{137}$-1 | *sdsN*$_{137}$*-1* under P$_{LlacO-1}$ control in pBRplac | SdsN$_{137}$-1 overproduction, from (2) |
| pDapZ | *dapZ* under P$_{LlacO-1}$ control in pBRplac | DapZ overproduction, this study |
| pCpxQ | *cpxQ* under P$_{LlacO-1}$ control in pBRplac | CpxQ overproduction, this study |

### Table S2: Oligonucleotides used in this study

| Name | Sequence | Use |
|---|---|---|
| **Strains construction** | | |
| 3'BaeRlacZ | TAACGCCAGGGTTTTCCCAGTCACGACGTTGTAAAACGACAACGATGCGGCAGGCGTCGGCTTCC | Construction Ptet-*baeR-lacZ* |
| 5'PtetBaeR | TAGAGATTGACATCCCTATCAGTGATAGAGATACTGAGCACACCGCTGGAACGGGATTTACAGAGAG | |
| 3'BasRlacZ | TAACGCCAGGGTTTTCCCAGTCACGACGTTGTAAAACGACGTTTTCCTCATTCGCGACCAGCATATAGC | Construction Ptet-*basR-lacZ* |
| 5'PtetBasR | TAGAGATTGACATCCCTATCAGTGATAGAGATACTGAGCACTTGCAGGAGAGTGAGTGAATGAAAATTC | |
| 3'EvgAlacZ | TAACGCCAGGGTTTTCCCAGTCACGACGTTGTAAAACGACGCCGATTTTGTTACGTTGTGCGAATGTG | Construction Ptet-*evgA-lacZ* |
| 5'PtetEvgA | TAGAGATTGACATCCCTATCAGTGATAGAGATACTGAGCACATACTTGTCGAATTATCTTAAAGGAAG | |
| 3'KdpElacZ | TAACGCCAGGGTTTTCCCAGTCACGACGTTGTAAAACGACAAGCATAAACCGATAGCCAATACC | Construction Ptet-*kdpE-lacZ* |
| 5'PtetKdpE | TAGAGATTGACATCCCTATCAGTGATAGAGATACTGAGCACGCAAACTGCCCCTGAACTTGAAG | |
| 3'NarPlacZ | TAACGCCAGGGTTTTCCCAGTCACGACGTTGTAAAACGACTTGTGCCCCGCGTTGTTGCAGGAAC | Construction Ptet-*narP-lacZ* |
| 5'PtetNarP | TAGAGATTGACATCCCTATCAGTGATAGAGATACTGAGCACTTTATAATGAAAATGATGCCAAAGC | |
| 3'PhoBlacZ | TAACGCCAGGGTTTTCCCAGTCACGACGTTGTAAAACGACAAAGCGGGTTGAAAAACGATATCCTGTACC | Construction Ptet-*phoB-lacZ* |
| 5'PtetPhoB | TAGAGATTGACATCCCTATCAGTGATAGAGATACTGAGCACGCATTAATGATCGCAACCTATTTATTAC | |
| 3'RstAlacZ | TAACGCCAGGGTTTTCCCAGTCACGACGTTGTAAAACGACTTCCCATGCATGAGGCGCAAAAAGATAGC | Construction Ptet-*rstA-lacZ* |
| 5'PtetRstA | TAGAGATTGACATCCCTATCAGTGATAGAGATACTGAGCACTTTTTATATCTACCGTGAATGTTATGAAC | |
| 3'UvrYlacZ | TAACGCCAGGGTTTTCCCAGTCACGACGTTGTAAAACGACCTGACTTGATAATGTCTCCGCATTAC | Construction Ptet-*uvrY-lacZ* |
| 5'PtetUvrY | TAGAGATTGACATCCCTATCAGTGATAGAGATACTGAGCACGACTAACTATCAGTAGCGTTATCC | |
| 3'YedWlacZ | TAACGCCAGGGTTTTCCCAGTCACGACGTTGTAAAACGACTTTTTTTACCGCTACGAATGAATAGC | Construction Ptet-*hprR-lacZ* |
| 5'PtetYedW | TAGAGATTGACATCCCTATCAGTGATAGAGATACTGAGCACATGAAGATTCTACTTATTGAAG | |
| 5'Ptet-ompR-35+30-lacZ | TAGAGATTGACATCCCTATCAGTGATAGAGATACTGAGCACTTACAAATTGTTGCGAACCTTTGGGAGT | Construction Ptet-*ompR-lacZ* |
| lacZ-37-73rev | GTTGGGTAACGCCAGGGTTTTCCCAGTCACGACGTTG | |
| narPmut2for | GATTCTAATAAAACGAGTGGAGACTACTATGCCTGAAGCAAC | *Construction narPmut2-lacZ* |
| narPmut2rev | GTTGCTTCAGGCATAGTAGTCTCCACTCGTTTTATTAGAATC | |
| rrnBt2-napF-for | GGCGCAGAAGGCCATCCTGACGGATGGCCTTTTTGCGTTTTTCCCGCCACTCTTTTGATC | *Construction PnapF-lacZ* |
| lacZ-17-napF-rev | CAGTGAATCCGTAATCATGGTCATAGCTGTTTCCTGTGTGAATTAATAAGCCATTTTTATAGC | |
| rrnBt2-PnarP_for | GGCGCAGAAGGCCATCCTGACGGATGGCCTTTTTGCGTTTGCGGAACGACAAAGAGAAAC | Construction P$_{narP-207+50}$-*lacZ* |
| PnarP+50-mut-lacZ_rev | GTCACGACGTTGTAAAACGACGGCCAGTGAATCCGTAATCATGTTCATAGCTGTTTCCTGTGTGAAGTGAAAACTACAACCTTG | |
| 5’Ptet-narP50_For | TAGAGATTGACATCCCTATCAGTGATAGAGATACTGAGCACTTGATTTTTTAGTTGTTTTTCTTGATGAG | 5’ primer for P$_{tet+1+50}$narP-lacZ construction |
| 40nt-rrnBt2-F | GAAGGCGAAGCGGCATGCATTTACGTTGACACCATCGAATGGCGCAGAAGGCCATCCTGA | When necessary, increase length of homology regions prior to |
| lacZ4-67rev | TAACGCCAGGGTTTTCCCAGTCACGACGTTGTAAAACGACGGCCAGTGAATCCGTAATCATGGT | |

| | | |
|---|---|---|
| | | recombineering into MG1508 |
| narP-FRT-kanR-FRT_for | GCCACCATTCTGTTCCTGCAACAACGCGGGGCACAATAATGTAGGCTGGAGCTGCTTCGAAG | Construction of *narP-linker-(3xFlag)-KanR* |
| linker-flag_for | GGTGCAGGTGCAGGTGCAGGTGCAGACTACAAAGACCATGACGGTGATTATAAAGAT | |
| narP-linker_for | GCCACCATTCTGTTCCTGCAACAACGCGGGGCACAAGGTGCAGGTGCAGGTGCAGGTGCA | |
| **Plasmids construction** | | |
| rprAmut2for | GATTTATAAGCATGGAAATCCCCACTCTGAAACAACGAATTGCTG | pRprAmut2 |
| rprAmut2rev | CAGCAATTCGTTGTTTCAGAGTGGGGATTTCCATGCTTATAAATC | |
| AatDapZ | GCTGACGTCAATATTTGTTATGGTGCAAAAATAAC | pDapZ |
| DapZEco | GCAGAATTCCAATTTAAAAACATAACACCAAAAATA | |
| AatCpxQ | CGAGACGTCTTTTCCTTGCCATAGACACC | pCpxQ |
| CpxQEco | GCTGAATTCGCAGCAGGCAAATTGAGGA | |
| EcoRI-RprA_rev | GCGAGAATTCGAAAGAGTGAGGGGCGAGGTAGC | pRprAproc |
| AatII_RprA60process_for | GCGTGACGTCATTGCTGTGTGTAGTCTTTGCCCATCT | |
| **Biotinylated probes for Northern-Blot** | | |
| RprA-probe | Bio-CAGGGGATTTCCATGCTTATAAATCAATATGTTGATTTATAACC | |
| RprAproc-probe | Bio-AAAAAAAGCCCATCGTGGGAGATGGGCAAAGACTA | |
| SsrA-probe | Bio-CGCCACTAACAAACTAGCCTGATTAAGTTTTAACGCTTCA | |
| ArcZ-probe | Bio-GGCTAGACCGGGGTGCGCGAATACTGCGCCAACACCAGGG | Same probe than ArcZNB4 in (6) |
